# Inactivation of the Niemann Pick C1 cholesterol transporter 1 (NPC1) restricts SARS-CoV-2 infection

**DOI:** 10.1101/2023.12.13.571570

**Authors:** Piergiorgio La Rosa, Jessica Tiberi, Enrico Palermo, Sofia Tiano, Mirko Cortese, John Hiscott, Maria Teresa Fiorenza

## Abstract

The Niemann Pick C1 (NPC1) protein is an intracellular cholesterol transporter located in the late endosome/lysosome (LE/Ly) and is involved in cholesterol mobilization. Loss-of-function mutations of the *NPC1* gene lead to accumulation of cholesterol and sphingolipids in LE/Ly, resulting in severe fatal NPC1 disease. Cellular alterations associated with NPC1 inactivation affect both the integrity of lipid rafts and the endocytic pathway. Because the angiotensin-converting enzyme 2 (ACE2) and type 2 serine transmembrane protease (TMPRSS2) of the SARS-CoV-2 Spike (S) protein also localize to lipid rafts, we sought to investigate the hypothesis that NPC1 inactivation would generate an intrinsically unfavorable barrier to SARS-CoV-2 entry. In this study, we demonstrate that NPC1 pharmacological inactivation or CRISP/R-Cas mediated ablation of NPC1 dramatically reduced SARS-CoV-2 infectivity. More specifically, our findings demonstrate that pharmacological inactivation of NPC1 results in massive accumulation of ACE2 in the autophagosomal/lysosomal compartment. A >40-fold decrease in virus titer indicates that this effectively prevents VSV-Spike-GFP infection by impeding virus binding and entry. A similarly marked decrease in viral infectivity is observed in cells that had NPC1 expression genetically abrogated. These observations were further confirmed in a *de novo* SARS-CoV-2 infection paradigm, where cells were infected with the naturally pathogenic SARS-CoV-2. Overall, this work offers strong evidence that NPC1 function is essential for successful SARS-CoV-2 infection, thus implicating NPC1 as a potential therapeutic target in COVID-19 management.

**Significance:** A significant convergence exists between the cellular alterations associated with NPC1 inactivation and the mechanistic processes of SARS-CoV-2 infectivity. These alterations affect the integrity of lipid-enriched plasma membrane microdomains and the endocytic pathway. Furthermore, the cholesterol-regulated ACE2 receptor protein that facilitates SARS-Cov-2 viral binding and entry is targeted to the autophagolysosomal compartment upon NPC1 inactivation, thus hindering virus-host cell interaction. To our knowledge, this study provides the first evidence that NPC1 function represents a crucial factor for SARS-CoV-2 infection and suggests therapeutic opportunities.

## Introduction

Cholesterol uptake occurs through the endocytic pathway and relies on the activity of the intracellular cholesterol transporter Niemann-Pick C1 (NPC1) to deliver cholesterol to various cellular compartments (1-3). NPC1 is a 13 transmembrane domain protein of the late endosome/lysosome (LE/Ly) limiting membrane involved in the egress of endocytosed cholesterol into the cytosol (3). Loss-of-function mutations in the *NPC1* gene cause endocytosed unesterified cholesterol and other sphingolipids to clog the LE/Ly compartment (4). The impairment of the endocytic sorting/degradation/recycling is responsible for a severe and ultimately fatal inherited disease with a systemic involvement, called Niemann-Pick type C (NPC) disease (5-7).

Because of entrapment within LE/Ly, normal amounts of cholesterol and sphingolipids fail to reach the plasma membrane (PM) and the endoplasmic reticulum (ER), where cholesterol homeostasis is fundamental to various signaling mechanisms (8, 9). For instance, at the level of PM, cholesterol is dynamically partitioned between the two fractions of accessible cholesterol and phospholipid- (mainly sphingomyelin) complexed cholesterol. Accessible cholesterol is metabolically active, that is readily used for covalent modification of cholesterol-regulated proteins (10), whereas sphingolipid-complexed cholesterol generates plasma membrane dynamic platform, named “rafts”. Here, cholesterol moieties ensure a tight packaging of fatty acid chains of sphingolipids, generating regions with a high structural order, to which various proteins are targeted (11,12).

The integrity of lipid rafts and the endocytic pathway is crucial for the initial steps of enveloped virus infection (13,14). The lipid envelope of SARS-CoV-2 is coated with the homotrimeric Spike (S) glycoprotein that mediates viral particle docking to the host cell receptor, angiotensin-converting enzyme 2 (ACE2) (15-17). After binding ACE2, the S protein is cleaved by the type 2 serine transmembrane protease, TMPRSS2, a membrane-bound enzyme that resides in close vicinity to ACE2, to mediate viral/plasma membrane fusion (18). TMPRSS2-mediated cleavage induces conformational changes in the S protein (referred to as “priming”), as well as in ACE2, constituting a crucial prerequisite for the endocytic entry of viral particles into the host cell (18, 19). Therefore, the host cell ACE2 receptor and the TMPRSS2 enzyme are involved in recognition and priming of the S protein, respectively (13), and their proper localization to lipid rafts is essential for virus entry (18). Alternatively, the S protein can undergo a series of cleavages by endo-lysosomal cathepsins B and L proteases when the virus is internalized *via* endocytosis (16, 20-22). This ensures the uncoating of the capsid shell and the subsequent release of the viral RNA genome into the cytosol for replication. The expression level of TMPRSS2 in different host cell types determines which mechanism prevails (21).

The requirement for NPC1 as an essential endosomal/lysosomal entry receptor is common among enveloped viruses, including Ebola, SARS, MERS and HIV viruses (23-26). Accordingly, NPC1 inactivation by imipramine or itraconazole has been shown to confer resistance to Ebola and HIV infection (27-29). The reliance of SARS-CoV-2 infection on the endocytic route suggested that inactivation of NPC1 could confer resistance to viral entry and propagation (30-33). However, underlying mechanisms are still unknown, with the exception of a recent study that identified NPC1 as an intracellular target of SARS-Cov-2 by demonstrating a physical interaction between SARS-Cov-2 nucleoprotein (N) and NPC1 (34).

In this study, we sought to evaluate whether transient inactivation of NPC1 would confer resistance to SARS-CoV-2 infection at an early post-entry step. To this end, we exploited human intestine (Caco-2) and lung (Calu-3) cells, and monkey kidney cells expressing human TMPRSS2 (VERO-76), to demonstrate that NPC1 inactivation interfered with SARS-CoV-2 infectivity. NPC1 inactivation was accomplished either by pharmacological inhibition with U18666A drug or by CRISP/R-Cas9 *NPC1* gene editing, and cell infection was carried out using a pseudotyped vesicular stomatitis virus (VSV) expressing SARS-CoV-2 S protein (35). These observations were further confirmed in a *de novo* SARS-CoV-2 infection paradigm, where cells were infected with the naturally pathogenic SARS-CoV-2. Overall, these findings demonstrate that the interference with NPC1 activity and cholesterol metabolism may represent a potential target to inhibit SARS-CoV-2 infectivity.

## Results

### Inactivation of NPC1 perturbs ACE2 and TMPRSS2 expression and localization

To evaluate the effect of NPC1 inactivation on SARS-CoV-2 infection, we employed the pharmacological inhibitor U18666A that blocks the cholesterol transport activity of NPC1 (36). The appropriate U18666A concentration and treatment duration for NPC1 inhibition were determined using dose-response curve analysis (Fig. 1A, Supplementary). The three cell types responded similarly to 24, 48 and 72 hours treatment with 0.02-200 uM U18666A concentrations (Fig. 1A, Supplementary). However, filipin staining revealed that a 24-hour treatment with 2 uM U18666A produced a robust NPC1 phenotype with minimal effect on cell survival (Fig. 1A, Supplementary, graphs). This observation prompted us to evaluate the mRNA and protein expression of NPC1, ACE2, and TMPRSS2 in VERO-76, Caco-2 and Calu-3 cells (37, 38) that were either untreated or treated for 24 hours with 2 μM U18666A (Fig. 1). Drug treatment modified transcript levels, as shown by a 2-fold increase in NPC1 transcript content in the three cell types. VERO-76 cells showed a similar 2-fold rise in ACE2 and TMPRSS2, but Caco-2 and Calu-3 cells showed about 2.5- and 1.5-fold decreases in ACE2 and TMPRSS2, respectively (Fig. 1A, B, C; see legend for statistical significance). In addition, the presence of U18666A significantly increased the expression of NPC1 protein in Calu-3 cells and decreased the expression of ACE2 in both Caco-2 and Calu-3 cells (Fig.1B, C).

**Figure 1.**
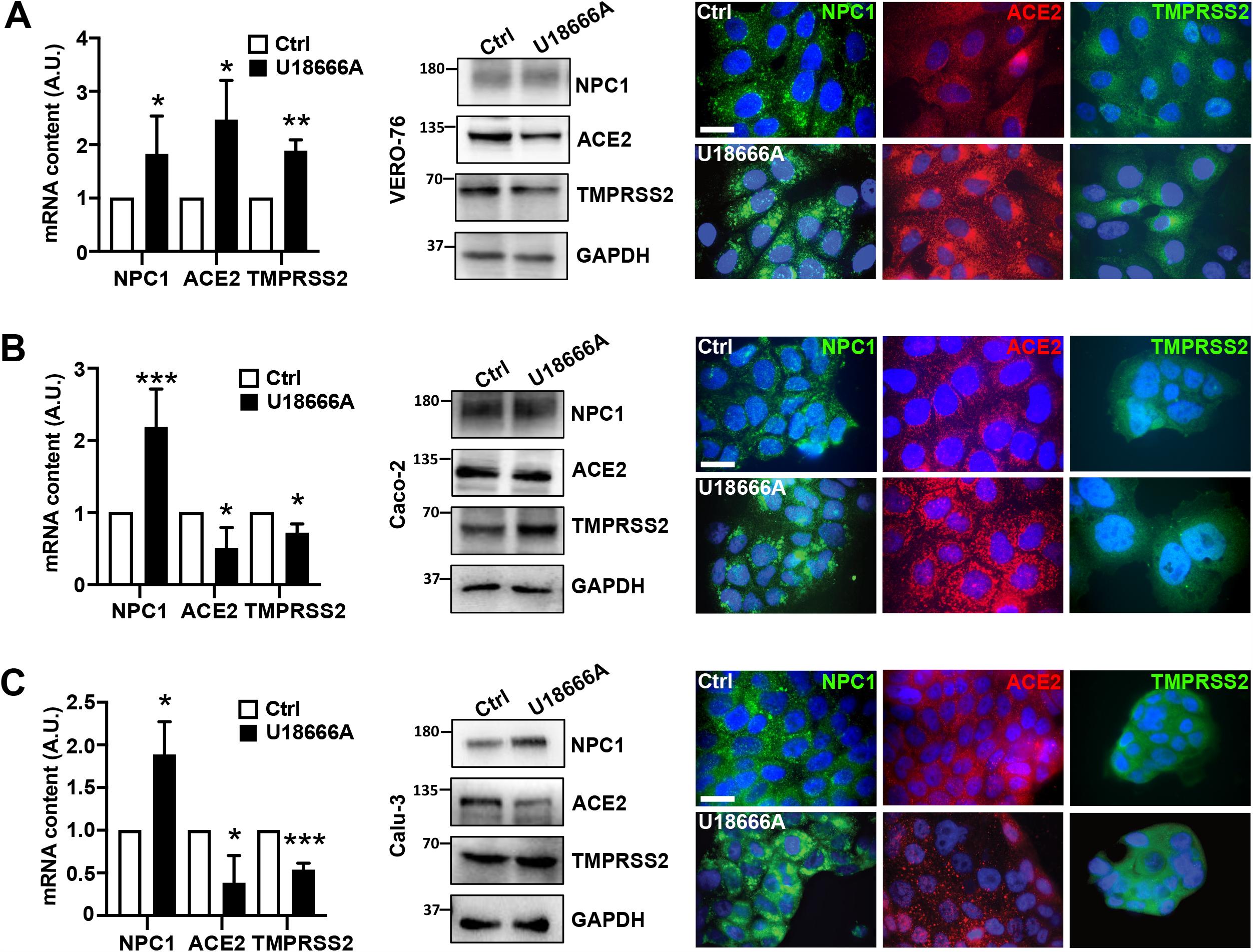
NPC1, ACE2 and TMPRSS2 expression patterns in untreated and U18666A-treated VERO-76, Caco-2 and Calu-3 cells. (A, B, C, left) Cells were treated with 2 μM U18666A or DMSO (vehicle, Ctrl) for 24 h. Total RNAs and proteins were extracted and analyzed by qRT-PCR and Western blot, respectively. Bars indicate relative abundance of NPC1, ACE2 and TMPRSS2 transcript levels normalized to RPL34 ribosomal protein RNA and expressed as fold-increase over control. Data are presented as mean ± SD of three independent experiments. *p < 0.05, **p < 0.005, ***p < 0.0005 vs Ctrl, calculated by Student’s t-test. Representative immunoblots of NPC1, ACE2 and TMPRSS2 protein content are shown (A, B, C). Densitometry of protein bands taking GAPDH as internal reference, was as follows: VERO-76 cells, NPC1, 1,66 ± 0,80; p= 0,28, n.s; ACE2, 0,84 ± 0,36; p= 0,44, n.s; TMPRSS2, 0,42 ± 0,22; p= 0,08, n.s. Caco-2 cells, NPC1, 1,38 ± 0,22; p= 0,59, n.s; ACE2, 0,59 ± 0,15; p= 0,027 *; TMPRSS2, 1,20 ± 0,09; p= 0,24, n.s.; Calu-3 cells, NPC1, 1,45 ± 0,19; p= 0,02 *; ACE2, 0,85 ± 0,08; p= 0,08, n.s.; TMPRSS2, 1,19 ± 0,07; p= 0,51, n.s. Representative images of immunofluorescence analysis of VERO-76 cells, Caco-2 and Calu-3 cells treated with 2 μM U18666A or DMSO (vehicle, Ctrl) for 24 h and processed for immunofluorescence with antibodies directed to NPC1, ACE2, and TMPRSS2, respectively. Nuclei were stained with Hoechst 33342. Images are representative of at least three independent experiments. Scale bar: 50 μM.

Next, we examined the effect of U18666A treatment on intracellular localization of NPC1 using immunofluorescence analysis. NPC1 acquired a prominent localization within the perinuclear organelles of all three cell types (Fig. 1A, B, C, immunofluorescence) (39), while a similar accumulation of ACE2 to the perinuclear region was observed in VERO-76 and Caco-2 cells, but less so in Calu-3. Similarly, VERO-76 showed a TMPRSS2 re-location pattern similar to ACE, but Caco-2 and Calu-3 cells showed no change in TMPRSS2 localization (Fig. 1A, B, C). These results indicate that drug treatment significantly changed the localization of NPC1 and ACE2 proteins, most likely as a result of the disruption of endocytic trafficking caused by NPC1 inactivation (39, 40-42).

### U18666A treatment targets NPC1 and ACE2 to the autophagolysosomal compartment

The observation that ACE2 acquired perinuclear organelle localization similar to NPC1 prompted us to characterize the compartment to which the two proteins were addressed, with particular reference to endocytic and/or autophagic networks (43, 44). Therefore, control and U18666A-treated Caco-2 cells were processed by double immunofluorescence analysis using antibodies directed against LAMP-2 and LC3B in order to detect the lysosomal and autophagosomal compartments, respectively (45). We observed a strong co-localization of NPC1-LC3B and NPC1-LAMP-2 within perinuclear dots (Fig. 2A, B), which is consistent with NPC1 targeting to the autophagosomal/lysosomal compartment. Remarkably, ACE2 was found to substantially co-localize with both LC3B and LAMP-2 (Fig. 2 C, D). VERO-76 treated with U18666A showed a similar pattern of NPC1 and ACE2 targeting to the autophagosomal/lysosomal compartment (Fig. 2S). Massive ACE2 accumulation in the autophagosomal/lysosomal compartment might prevent virus-host cell interaction, reducing the susceptibility of U18666A-treated cells to SARS-CoV-2 infection.

**Figure 2.**
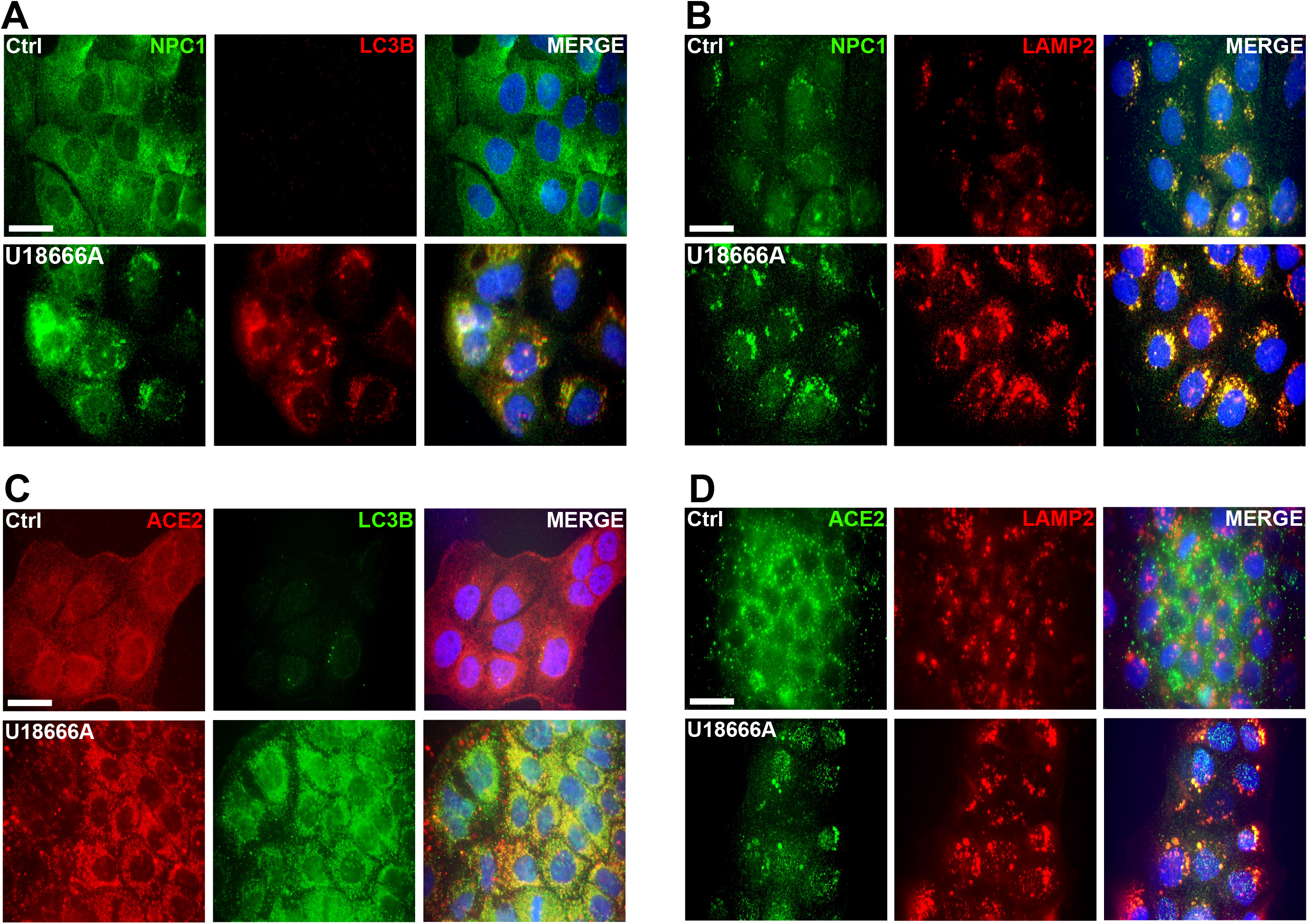
Characterizing how cell treatment with U18666A drug affects the intracellular localization of NPC1 and ACE2. (A-D) Representative images of immunofluorescence analysis of Caco-2 cells treated with 2 μM U18666A or DMSO (vehicle, Ctrl) for 24 h and then subjected to double immunofluorescence experiments by the incubation with the following pairs of antibodies: α-NPC1 (green), α-LC3B (red) (A), α-NPC1 (green), α-LAMP2 (red) (B); α-ACE2 (red), α-LC3B (green) (C), α-ACE2 (green), α-LC3B (red) (D). Nuclei were stained with Hoechst 33342. Images are representative of at least three independent experiments. Scale bar: 50 μM.

### *NPC1 inactivation confers resistance to* SARS-CoV-2 Spike pseudotyped virus *entry and propagation*

To directly investigate the possibility that cell treatment with U18886A interfered with the docking of SARS-CoV-2 Spike protein to the plasma membrane of host cells, U18666A- or mock-treated Caco-2 cells were infected with pseudotyped VSV-Spike-GFP virus (35), and infectivity measured by GFP expression. U18666A treatment dramatically reduced GFP fluorescence by >90%, essentially blocking VSV-Spike-GFP infection (Fig. 3A). Consistent with these results, virus titers were reduced by >40-fold, compared to untreated infected cells (Fig. 3B). The percentage of apoptotic cells in U18666A-treated Caco-2 cells was comparable to that in control and U18666A-treated mock cells (Fig. 3C), providing additional evidence of the resistance to virus infection determined by NPC1 inactivation. In contrast, U18666A-untreated infected cells (Fig. 3C) showed a dramatic VSV-induced cytopathic effect, with approximately 30% apoptotic cell death. A lower concentration of U18666A (0.2 uM, 72 hours, see dose-response curves of Fig. 1A, Supplementary) produced results that were comparable (not shown). NPC1 inactivation in Calu-3 and VERO-76 cells produced similar patterns of resistance to VSV-Spike-GFP infection and propagation (Fig. 3 A-F, Supplementary). Decreased vulnerability to VSV-Spike infection was consistent with the marked decrease in Spike expression in VERO-76 cells treated with U18666A when compared to control (p ≤0.0001) (Fig. 3G, Supplementary). Lastly, we assessed the virus ability to bind to the ACE2 receptor and entry (Fig. 3D), observing that NPC1 inactivation results in a considerable reduction in both viral binding and subsequent entrance as evidenced by the 30% decrease in binding and the 50% decrease in viral RNA and GFP expression seen in cells treated with U18666A (Fig. 3D, bars).

**Figure 3.**
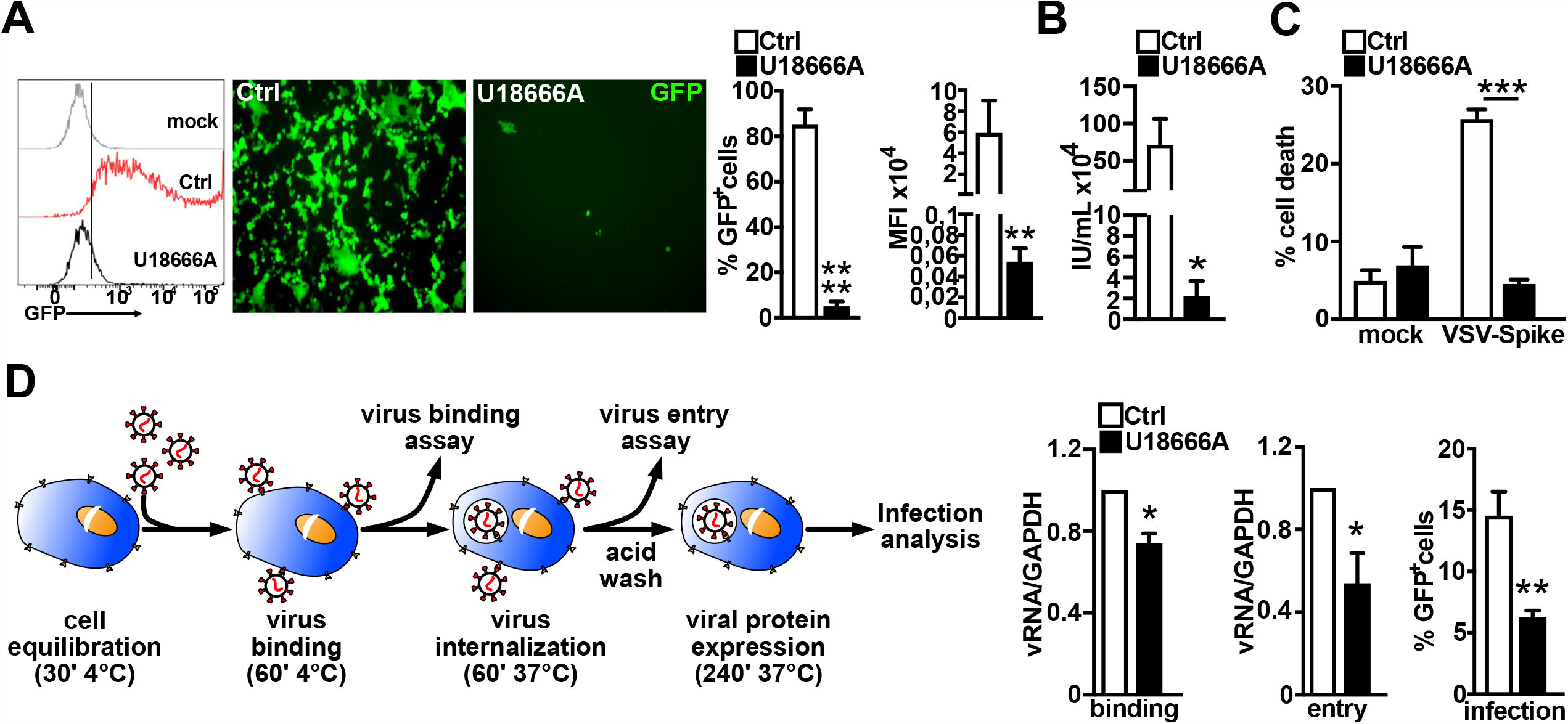
The inhibition of NPC1 counteracts VSV-Spike entry. (A) Cells were seeded in a 12-well plate and treated with either 2 uM U18666A inhibitor or DMSO (vehicle, Ctrl) for 24 h. Cells were then infected with VSV-Spike-GFP (MOI 0.1), harvested 24 h after infection and analyzed for GFP expression. An example of a FACS analysis’s output showing the percentage of GFP-positive cells/experimental/group. Representative images of immunofluorescence analysis of VSV-Spike-GFP infected cells and quantitative analysis of GFP-positive cells (histograms). **p< 0.0001; ****p < 0.0001. (B) Control and U18666A-treated cells were infected with VSV-Spike-GFP (MOI 0.1). Supernatants were collected after 24 h and used to infect Vero E6 cells; at 7h post-infection the fraction of GFP-positive cells was quantified by flow cytometry and viral titer, expressed as infection units (IU), was determined. Histograms represent mean ± SD from three independent experiments. *p < 0.0001. (C) Cells were harvested 24 h following VSV-Spike-GFP (MOI 0.1) infection and stained for 7-AAD to quantify the fraction of dead cells by flow cytometry. Histograms represent the mean ± SD from three independent experiments. ***p < 0.0001. (D) The effect of NPC1 inhibition on VSV-Spike-GFP binding to cell membrane and cell entry. A scheme of the experimental paradigm. A pre-incubation at 4 °C preceded Caco-2 cell infection with SARS-CoV-2 Spike pseudotyped virus (MOI 1) and was followed by 1 h cell incubation at the same temperature. Cells were then harvested and the amount of pseudoviral particles bound to cell membrane was determined by qPCR of genomic virus RNA. Alternatively, cells were transferred to a temperature of 37 °C for 1 h and then processed for qPCR of genomic virus RNA. Histograms indicate the content of viral genome in Ctrl and U18666A-treated cells determined after “virus binding” and after “virus entry”, respectively. GAPDH was taken as reference for normalization. Data are presented as mean ± SD from three independent experiments. *p < 0.0001. Right panel shows the percentage of GFP-positive cells measured by flow cytometry at 4h post-infection. Data are presented as mean ± SD from one experiment performed in triplicate **p < 0.0001.

### NPC1 loss-of-function restricts SARS-CoV-2 infection

The ability of *NPC1* loss to provide resistance to VSV-Spike-GFP entry was then evaluated in *NPC1*^*-/-*^ Caco-2 cells, generated by CRISP/R-Cas9 gene editing. Filipin staining revealed two clones, named LG5 and LD6, that had the intracellular cholesterol accumulation characteristic of NPC1-deficient cells (Fig. 4A, Supplementary). Consistently, NPC1 protein content was found to be below detection in LG5 cells and barely detected in LD6 cells (Fig. 4A, Supplementary). Moreover, residual NPC1 protein of LD6 cells appeared to be targeted to the autophagosome/lysosome compartment (Fig. 4B, Supplementary). No *NPC1* transcript expression was detected in LG5 with primers specific for the targeted exon 4 and a significant reduction of amplification compared to control cells with primers downstream from exon 4 (Figure 4C, Supplementary). A slight reduction of *NPC1* transcript amplification in LD6 compared to control cells was also observed.

**Figure 4.**
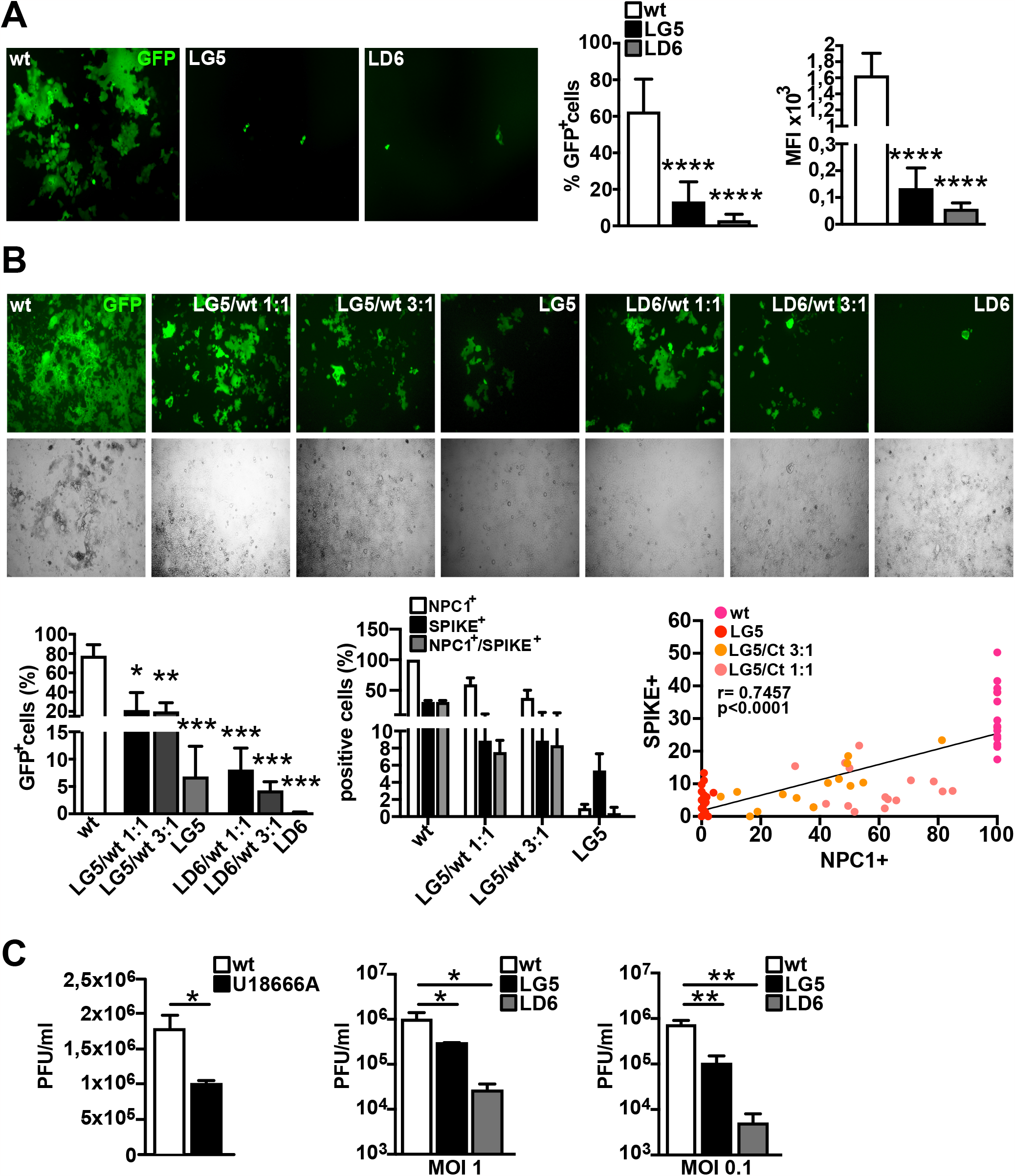
NPC1 loss-of-function counteracts VSV-Spike-GFP and SARS-CoV-2 infection. (A) Cells, either wild-type (wt) or *NPC1*-edited, were infected with VSV-Spike-GFP (MOI 0.1) and pseudoviral infectivity was measured 24 h post-infection by determining GFP expression by immunofluorescence and flow cytometry. Histograms represent the mean ± SD from three independent experiments. (B) LG5 and LD6 clone cells alone, or mixed to wt cells in two different ratios, *i*.*e*. 1:1 or 3:1, were infected with VSV-Spike-GFP (MOI 0.1) and pseudoviral infectivity was measured 24 hr post-infection by determining the fraction of GFP-positive cells (B, bars on the left). A separate set of LG5 infected cells was processed for NPC1/Spike double immunofluorescence to determine the relative fraction of NPC1-positive, Spike-positive and NPC1/Spike-positive cells out of total cells (B, bars in the middle). Pearson’s correlation coefficient between NPC1 and Spike expression (B, graph on the right). (C) Untreated and U18666A-treated cells were infected with native SARS-CoV-2 virus and released infectivity was determined from supernatants collected 24 h post-infection (C, bars on the left). Wt, LG5 and LD6 cells were infected with SARS-CoV-2 at either 1 or 0.1 MOI and released infectivity was determined from supernatants collected 24 h post infection (C, middle and right bars). Histograms represent the mean ± SD from three independent experiments. *, p ≤ 0.05; **, p ≤ 0.05; ***, p ≤ 0.05; ***, p ≤ 0.05.

Consistent with the solid NPC1 phenotype displayed by LG5 and LD6 cells (Fig. 4A, Supplementary), cells of both clones were resistant to SARS-CoV2 infection when challenged with VSV-Spike-GFP; LG5 and LD6 displayed ∼13% and ∼3% GFP positive cells respectively, *vs* ∼60% GFP positive cells in *wild-type* (*wt*) Caco-2 cells (Fig. 4A, left). As an additional proof of the strict dependency of virus infection on the NPC1 expression, we mixed LG5 or LD6 cells with *wt* Caco-2 cells at different ratios, and observed that the increase in the fraction of LG5 or LD6 cells was accompanied by a decrease of GFP-positivity (Fig. 4B). The relationship between NPC1 expression and virus propagation was further supported by the direct correlation observed between the fraction of Spike- and NPC1-expressing cells within the mixed population (r 07457; p 0.0001; Fig. 4B).

The role of NPC1 as a critical viral restriction factor was further corroborated by infecting untreated and U18666A-treated Caco-2 cells with pathogenic SARS-CoV-2 (BAVPAT-1) (Fig. 4C, left bars). SARS-CoV-2 infectivity was reduced by a 42% in U18666A-treated cells compared to control, as determined by virus titration (Fig. 4C, bars on the right). A dramatic reduction of SARS-CoV-2 infectivity was also observed when *wt* and *NPC1*^*-/-*^ Caco-2 cells, respectively, were infected with pathogenic SARS-CoV-2. Specifically, SARS-CoV-2 infectivity was reduced by a 70% (LG5) and 97% (LD6) at MOI 1; when cells were infected at MOI 0.1, SARS-CoV-2 infectivity showed a further drop, with a 86% and 99% reduction in LG5 and LD6 cells, respectively (Fig. 4C, middle and right).

To gain a mechanistic insight on the resistance of LG5 and LD6 clone cells to SARS-CoV-2 infectivity, the expression patterns of ACE2 and TMPRSS2 were determined in LG5 and LD6 clones. Compared to *wt* Caco-2 cells, a 50% reduction in ACE2 protein expression was detected in both LG5 and LD6; expression of TMPRSS2a was also reduced, although these values did not reach statistical significance (Fig. 5 Supplementary). Likewise, ACE2 transcript levels were reduced in both LG5 and LD6 cells, whereas TMPRSS2 transcripts decreased only in LD6 cells (Fig. 5, Supplementary). A pronounced co-localization of ACE2 with LC3B and LAMP2 was also detected in both clones (Fig. 5, Supplementary), which recapitulates the expression patterns observed in U18666A-treated cells. Overall, these results indicate that the internalization of ACE2 as a consequence of NPC1 ablation interferes with the docking of SARS-CoV2 Spike protein to the plasma membrane, thus increasing resistance of Caco-2 cells to SARS-CoV-2 entry.

## Discussion

The lipid microenvironment of various cellular organelles is known to be a critical factor for the infectivity of enveloped viruses, including SARS-CoV-2 (22, 46-48). For instance, the depletion of cholesterol and sphingolipids at the plasma membrane (PM) hinders virus entry by interfering with the docking of the Spike protein to the ACE2 receptor and the subsequent TMPRSS2 priming or clatrin- and caveoale-dependent endocytosis (13,18). The disturbance of nonvesicular lipid transport *via* membrane contact sites (MCVs), with particular reference to LE/L-ER MCVs, hampers virus replication, because double membrane vesicles (DMVs) that act as platform for viral replication originate from cholesterol-rich ER sub-compartments (49). With the present study, we demonstrate that pharmacological or genetic inactivation of NPC1 interferes with SARS-Cov-2 binding and entry, highlighting the fact that dysregulation of intracellular cholesterol trafficking and biosynthesis modulates the infectivity of SARS-CoV-2, as well as several viruses (23, 50-53).

U18666A has been widely used to model Niemann Pick C disease (36, 54-57) and to investigate the effect of NPC1 inhibition on viral replication (23,28, 58-61), whereas studies investigating its effect on viral binding and entry are scarce. The various cell types of this study had a comparable response to U18666A treatment exhibiting the expected phenotype of intracellular cholesterol accumulation, an increase in *NPC1* mRNA expression and NPC1 protein targeting to perinuclear organelles (reviewed in 43). On the other hand, our finding that U18666A-dependent NPC1 inhibition robustly alters the intracellular localization of ACE2 protein is a novel finding, which indicates a modification of ACE2 trafficking and/or partitioning among cell compartments. Specifically, the accumulation of ACE2 within the autophagolysosomal compartment is consistent with a dysfunctional autophagosome-lysosome fusion brought on by U18666A-dependent NPC1 inhibition (62).

Concerning the change in ACE2 intracellular localization, we take into account two possible scenarios: either newly synthesized ACE2 protein is prevented from moving to the PM by a malfunctioning ER-Golgi secretory pathway (62), or reduced cholesterol levels at the PM cause ACE2 internalization by upsetting the ideal microenvironment for its activity (47, 63). Both scenarios are consistent with the observation that the low concentrations of U18666A employed in this study decreases the delivery of cholesterol from lysosomes to both PM and ER (64).

Although more research is needed to fully understand the effects of NPC1 inhibition on ACE2 portioning inside PM and subsequent fate, its accumulation in the autophagolysosomal compartment indirectly indicates that treatment of cells with U18666A reduces the amount of ACE2 fraction available at the PM. This offers a clear explanation for the high resistance of cells treated with U18666A to SARS-CoV-2 infection. In fact, it is well known that cholesterol-regulated ACE2 localization to suitable PM lipid clusters promotes viral binding and entrance, both of which - according to our data - are markedly reduced after NPC1 inhibition. Therefore, we can presume that cell treatment with U18666A hinders SARS-CoV-2 infection by modifying the cholesterol-dependent ACE2 association to PM lipid microenvironment required for virus binding (63). This conclusion is in agreement with the study by Takano et al., showing that membrane cholesterol levels are critical for the successful infection of host cells by feline coronavirus (60). Additionally, a recent study has demonstrated that SARS-CoV-2 infectivity is strictly dependent on ACE2 portioning to appropriate PM lipid clusters (65). Conversely, because lysosomal pH is barely affected by the relatively low concentration of U18666A (2 μM) that has been used in this study to achieve NPC1 inhibition, the possibility of an antiviral effect based on the block of cathepsin protease action by an increase in lysosomal pH seems unlikely (66).

In addition to the impact on viral binding and entry, our results show a noteworthy reduction in released infectivity in U18666A-treated cells. Although this is in line with an impairment of viral replication as discussed above, the possibility exists that the egress of virions from infected cells is also impaired. This is because SARS-CoV-2 exploits lysosomal trafficking instead of the secretory pathway for egress (67). Consequently, it is expected that the disruption of lysosomal transport linked to NPC1 inactivation (68) will impact the release of viral particles.

Further validation of NPC1’s crucial function in controlling cell vulnerability to SARS-CoV-2 infection is provided by experiments conducted in Caco-2 cells that have had their NPC1 expression abrogated via CRISP/R-Cas9. The two generated clones (LG5 and LD6) showed a significantly high resistance to the infection when challenged with VSV-Spike-GFP under conditions comparable to those used in U18666A-treated cells. We also provide a more concrete demonstration of NPC1 loss-of-function as a defense against SARS-CoV-2 by infecting increasing amount of LG5 and LD6 cells mixed to a fixed amount of wild-type (wt) cells. Using this approach, we were able to confirm that NPC1 expression and infection ratios are strictly correlated. Indeed, the identification of infected cells by means of either GFP or Spike expression provided the opportunity to analyze immunofluorescence data for Pearson correlation coefficient, observing the existence of a significant linear correlation between NPC1-expressing cells and Spike-positive cells. Lastly, our findings further support the study hypothesis, showing that when challenged with natural SARS-CoV-2, either U18666A-treated Caco-2 cells or LG5/LD6 cells showed a significant reduction of infection rates compared to control or wild-type cells, respectively. Moreover, the intracellular localization patterns of ACE2, in conjunction with immunofluorescence analyses of LAMP-2 and LC3B, indicate that the molecular basis of infection resistance is similar in U18666A-treated and NPC1-KO cells.

Overall, this study provides compelling evidence that NPC1 function is an important component of substantial SARS-CoV-2 infection and that blocking it may have significant therapeutic benefits.

## Materials and Methods

### Cell Culture and treatments

African green monkey kidney Vero-E6 and VERO-76 (hTMPRSS2) cells (RRID: CVCL_0603), Intestinal epithelial cell line Caco-2 (RRID: CVCL_0025), and human airway epithelial Calu-3 cells (RRID: CVCL_0609), were cultured in Dulbecco’s modified Eagle’s medium (DMEM, Merck, cat #D5671) supplemented with 10% fetal bovine serum (FBS, Merck, cat #F2442), 100 U/mL penicillin/streptomycin (Merck, cat #P4333), sodium pyruvate (Merck, cat #S8636), MEM non-essential Amino Acid (Merck, cat #M7145) and Normocin (Invivogen, cat #ant-nr1), in a humidified incubator with 5% CO_2_ at 37 °C. The NPC1 inhibitor U18666A drug was purchased from Merck (cat #U3633) and dissolved in DMSO, obtaining a 2 mM stock solution.

### Filipin staining and Immunofluorescence

Cells were fixed with 4% (v/v) PFA (Merck, cat #F8775) for 15 min, washed thrice with Dulbecco’s Phosphate Buffered Saline (D-PBS, Euroclone, cat #ECB4004L). For filipin staining, cells were incubated with filipin complex (Merck, cat #F-9765; 0.15 mg/ML in D-PBS) for 2 h and after several washes in D-PBS were visualized by fluorescence microscopy. For immunofluorescence assays, fixed cells were permeabilized with 0.1% Triton X-100 (Merck, cat #T8787) in D-PBS supplemented with 1% BSA (Merck, cat #A2153) and incubated overnight at 4° C with the following primary antibodies: α-NPC1 (1:100, cat #ab134113, Abcam), α-ACE2 (1:200, cat #66699-1-Ig, Proteintech), α-TMPRSS2 (1:200, cat #14437-1-AP, Proteintech), α-SPIKE (1:100, cat#703958, Invitrogen), α-LC3-B (1:100, cat #GTX632501, GeneTex), α-LAMP2 (1:100, cat #MA1-205, Thermofisher). Secondary antibody were Alexa Fluor 488 anti-rabbit IgG (Invitrogen, cat #A11070) and Alexa Fluor anti-mouse IgG (Invitrogen, cat #A32727), both used at 1:500 dilution for 1 hr incubation at RT. Nuclear staining was performed with Hoechst (Merck, cat #94403) for 15 min. Slides were mounted using Prolong Gold mounting solution (Invitrogen, cat #00-4958-02). Randomly selected fields for each sample were acquired using a DFC3000 G inverted microscope (LEICA Geosystems).

### Immunoblotting

Cells were lysed in a buffer containing: 50 mM Tris-HCl pH 7.6, 150 mM NaCl, 2 mM EDTA, 1% NP-40, 0,5% Triton X-100, supplemented with protease inhibitor cocktail (SERVA, cat #39102.01). After protein separation by SDS-PAGE and transfer to polyvinyl difluoride (PVDF) membranes (Amersham, United Kingdom), blots were incubated overnight at 4°C with the following antibodies: α-ACE2 (1:1000), α-TMPRSS2 (1:2000), α-NPC1 (1:2000), α-GAPDH (1:5000, cat #E-AB-20059, Elabscience) in TBS containing 0.1% Tween-20 and 5% BSA. Protein bands were detected by enhanced chemiluminescence (Advansta, cat #K-12043-D10) and visualized using iBright 1500 Imaging System (Invitrogen, cat #A43678).

### RNA isolation and qRT-PCR

Total RNA was extracted using Total RNA purification plus kit (Norgen Biotek Corp., cat #48400) acc. to manufacturer instructions. 1 μg of RNA was retrotranscribed using OneScript Plus cDNA Synthesis Kit (abm, cat #G236). cDNAs were amplified in qPCR assays, using 2x SensiFAST SYBR Lo-ROX Mix (Bioline, cat #BIO-94002) following manufacturers’ instructions. All primers used in RT-PCR and qPCR experiments are reported in Supplementary table 1. Ribosomal Protein L34 (RPL34) mRNA was used for normalization in qPCR experiments.

### NPC1 gene editing

Caco-2 cells were co-transfected with eSpCas9-GFP protein (Merck, cat #ECAS9GFPPR) and sygRNA (Merck, cat #HSPD0000028649) targeting the 4^th^ exon in the *NPC1* gene, using Lipofectamine 2000 (Invitrogen, cat #11668-030) as per manufacturer’s instructions. Cells were harvested 24 h after transfection, and the top EGFP-expressing cells were sorted into 96-well plates as single clones and as enriched EGFP-expressing population (100 EGFP positive cells per well) depending on the experiment. To screen for clones with *NPC1* gene disruption, genomic DNA was analyzed by PCR. NPC1 protein depletion and NPC1 gene disruption were confirmed by Western blot and qPCR.

### Pseudoviral particles titration and cell infection

VSV-Spike-GFP - a replication-competent VSV expressing eGFP in the first position of the genome as well as a modified version of the SARS-CoV-2 spike in place of the native VSV glycoprotein - was a kind gift of Prof. David Olagnier (Aarhus University, Aarhus C 8000, Denmark) and was originally obtained from Prof. Paul W. Rothlauf as described (35). VSV-Spike-GFP was propagated in VERO-76-hTMPRSS2 cells; briefly, cells were infected with VSV-Spike-GFP at MOI of 0.01 for 48 h, supernatant was collected, centrifuged 5’ at 300 x g and then filtered using a 0.22 μm bottle-top vacuum filter. Virus was concentrated by ultracentrifugation at 18,000 x g for 90’ at 4 °C and then purified on a 20% sucrose cushion at 135,000 x g for 90’ at 4 °C using a Beckman SW32Ti Swing Bucket Rotor. Purified virus was resuspended in PBS and virus titer was quantified by flow cytometry analysis of GFP expression in Vero E6 cells (% GFP positive cells*#infected cells/mL of virus), as previously described (69). Titers were expressed as IU/mL. In infection experiments, cells were pre-treated with U18666A inhibitor or DMSO (vehicle, Ctrl) for 24 h and then were incubated with virus (MOI 0.1) in serum-free medium for 1 h in a humidified incubator with 5% CO_2_ at 37 °C; after incubation, the virus containing medium was removed and replaced with fresh complete medium for 24 h prior to analysis by FACS or immunofluorescence.

In experiments that determined the virus ability to bind to the ACE2 receptor and entry, a 30 min pre-incubation at 4 °C preceded cell infection and was followed by 1 h cell incubation at the same temperature. Cells where then harvested and the amount of pseudoviral particles bound to cell membrane was determined by qPCR of viral RNA. Alternatively, cells were transferred to a temperature of 37 °C for 1 h and then processed for qPCR of genomic virus RNA. Total RNA was isolated by column separation using the RNeasy Kit (Qiagen) acc. to manufacturer’s instructions and the concentration was measured with Nanodrop 2000 (Thermo Fisher Scientific). A quantity of RNA in the range 200 – 500 ng was used for cDNA synthesis using the PrimeScript RT-PCR Kit (Takara-Bio). Quantitative PCR was then performed using Taqman Fast Advanced MasterMix with Universal Probe Library Probes (Roche) with specific primers designed using the Roche Lifescience Assay Design Center (https://lifescience.roche.com/en_it/brands/universal-probe-library.html#assay-design-center) on a StepOnePlus Real-Time PCR System (Thermo Fischer Scientific). A relative quantification method was used, with GAPDH as housekeeping gene. Primers used: VSV-N: Fw-CCTGATGACATTGAGTATACATCTCTT, Rev-GGATCCTACTGCATAAGCGTACA; GAPDH: Fw-AGCCACATCGCTCAGACA, Rev-GCCCAATACGACCAAATCC.

### Plaque assay

To compute the number of PFU/mL in the supernatants collected from infected Caco-2 clones, 2.5 x 10^5^ Vero E6 cells were seeded in 24 well plates. The day after seeding, 10-fold dilutions of the supernatants were added to the Vero E6 cells and infected cells were incubated at 37 °C for 2 hours. After that, the inoculum was removed and replaced with 1 mL of plaquing medium 0.9% carboxymethyl-cellulose (Sigma-Aldrich) in MEM. After a 72 h incubation, Vero E6 cells were fixed in 5% formaldehyde for 30 min and subsequently transferred in a 6% formaldehyde bath for 30 minutes for inactivation before being transported outside the BSL-3 area. Plates were then rinsed with tap water and incubated for 15 minutes with 1% Crystal Violet (Sigma-Aldrich), rinsed again and PFU/mL were counted on stained monolayers.

### SARS-CoV-2 infection on Caco-2 clones

The day before the infection, 5 x 10^4^ cells of each Caco-2 clone were seeded in a 24 well plate. 24 h after seeding, cells were infected in technical duplicates with SARS-CoV-2 (BAVPAT-1) at MOI 1 and 0.1. After 2 h the viral inoculum was removed, cells were washed twice in PBS and DMEM with 2% FBS was added on top. Supernatants were collected at 24 h post-infection and viral titer was assessed by plaque assay.

### SARS-CoV-2 infection in presence of U18666A inhibitor

Wild-type Caco-2 cells were seeded in a 24 well plate at 5 x 10^4^ cells per well and after 24 h were treated either with DMSO or 2 uM U18666A. After 24 h of treatment, medium was replaced with DMEM with 2% FBS without compound and cells were infected with SARS-CoV-2 at MOI 0.1. Viral inoculi were removed after 2 h, cells were washed twice in PBS and DMEM with 2% FBS was added to each well. Supernantants were collected at 24 h post-infection and viral titers were quantified by plaque assays on Vero E6 cells.

## Acknowledgments

We thank Giovanna Peruzzi for support with FACS experiments and Robert P Erickson for critical reading of manuscript. M.C. is supported by the Human Technopole Early Career Fellowship Programme (HT-ECF 763 Programme). The COVID19 Special Grant - GSP20006_Covid050 from the Telethon Foundation and RG1221816B9646F5 – Sapienza University – to MTF provided funding for this work.

