## Supplementary materials for "Inactivation of the Niemann Pick C1 cholesterol transporter 1 (NPC1) restricts SARS-CoV-2 infection"

##### Supplementary figure legends

**Figure 1S**. Validation of NPC1 phenotype obtained by U18666A drug administration in cell lines originating from various tissues. (A) Vero E6-TMPRSS2, Caco-2 and Calu-3 cells were treated with increasing concentration of U18666A (0.02 µm-20 µm) for 24 h and stained with filipin; control cells were treated with 0.2% DMSO (vehicle, Ctrl). Cell growth analysis at 24, 48, and 72 hr following treatment initiation with increasing U18666A concentrations (0.02 µM-200 µM). (B) Vero E6-TMPRSS2, Caco-2 and Calu-3 cells were treated either with 0.2 µM or 2 µM U18666A for 24, 48, and 72 hr and imaged following filipin staining. Representative images of three independent experiments are shown in A and B. Scale bar: 50 µM. $^{**}p < 0.01$, $^{****}p < 0.0001$ *vs* Ctrl, calculated by TWO WAY ANOVA.

**Figure 2S.** The intracellular localization of NPC1 and ACE2 in VERO-76 cells varies upon U18666A drug treatment. Representative images of immunofluorescence analysis of cells treated with 2 µM U18666A or DMSO (vehicle, Ctrl) for 24 h and then subjected to double immunofluorescence by the incubation with the following pairs of antibodies: α-NPC1 (green), α-LC3B (red) (A), α-NPC1 (green), α-LAMP2 (red) (B); α-ACE2 (red), α-LC3B (green) (C), α-ACE2 (green), α-LC3B (red) (D). Nuclei were stained with Hoechst 33342. Images are representative of at least three independent experiments Scale bar: 50 µM.

**Figure 3S.** The inactivation of NPC1 in Calu-3 and VERO-76 cells counteracts VSV-Spike entry. (A) Calu-3 cells were treated with U1866A (2 uM) or DMSO (vehicle, Ctrl) for 24 hr. Cells were then infected with VSV-Spike-GFP (MOI 0.1), harvested 24 h after infection and analyzed for GFP expression. A representative output of FACS analysis displaying the fraction of infected, GFP-positive cells/experimental group (A left). Representative images of fluorescence microscopy analysis of VSV-Spike-GFP infected cells and quantitative analysis of GFP-positive cells (bars). $^{***}p < 0.0001$. (B) Control and U18666A-treated cells were infected with VSV-Spike-GFP (MOI 0.1). Supernatants were collected after 24 h and used to infect Vero E6 cells; at 7 h post-infection the fraction of GFP-positive cells was quantified by flow cytometry and viral titer, expressed as infection units (IU), was determined. Histograms represent mean ± SD from three independent experiments. *p < 0.0001. (C) Cells were harvested 24 h following VSV-Spike-GFP (MOI 0.1) infection and stained for 7-AAD to quantify the fraction of dead cells by flow cytometry. Histograms represent the mean ± SD from three independent experiments; n.s., non significant. (D) VERO-76 cells were treated with 2 uM U1866A inhibitor or DMSO (vehicle, Ctrl) for 24 h. Cells were then infected with VSV-Spike-GFP (MOI 0.1), harvested 24 h after infection and analyzed for GFP expression by flow cytometry. A representative output of FACS analysis displaying the fraction of infected, GFP-positive cells/experimental group (D left). Representative images of immunofluorescence analysis of VSV-Spike-GFP infected cells and quantitative analysis of GFP-positive cells (bars). *p < 0.0001. (E) Control and U18666A-treated cells were infected with VSV-Spike-GFP (MOI 0.1). Supernatants were collected after 24 h and used to infect VERO-76 cells; at 7 h post-infection the fraction of GFP-positive cells was quantified by flow cytometry and viral titer, expressed as infection units (IU), was determined. Histograms represent mean ± SD from three independent experiments. *p < 0.0001. (F) Cells were harvested 24 h following VSV-Spike-GFP (MOI 0.1) infection and stained for 7-AAD to quantify the fraction of dead cells by flow cytometry. Histograms represent the mean ± SD from three independent experiments; *p < 0.0001. (G) Representative images of fluorescence microscopy analysis of VSV-Spike-GFP infected cells processed by immunofluorescence with α-Spike antibodies. Histograms represent the fraction of Spike-positive cells. **** p ≤0.0001. Scale bar: 50 μM.

**Figure 4S.** Validation of NPC1 loss-of-function obtained by CRISPR/Cas9 gene editing. (A) Caco-2 cells were edited using a RNA-guided Cas9 nuclease targeting exon 4. Two clones, labeled as LG5 and LD6, nicely mimicked the typical intracellular cholesterol accumulation of NPC1 phenotype, as detected by filipin staining. Representative immunoblot and quantitative analysis (bars) showing that NPC1 protein is below detection in LG5 and barely detected in LD6 clone cells, compared to wt cells. Data are presented as mean ± SD of three independent experiments. * p ≤ 0.05; ** p ≤ 0.005. (B) Characterization of LG5 and LD6 clone cells by immunofluorescence analysis with the following pairs of antibodies: α-NPC1 (green), α-LC3B (red) (B, left), α-NPC1 (green), α-LAMP2 (red) (B, right). Merged images are also shown. (C) A scheme of primer design to assess NPC1 gene editing by transcript amplification. Four primer pairs were designed to encompass nucleotide sequences, as follows: 1) exons 2 and 3; 2) exon 4; 3) exons 4 and 5; 4) exons 5 and 6 (C, scheme). Bars indicate the relative abundance of RT-PCR amplified products from LG5 and LD6 clones with the various primer pairs, as indicated.

**Figure 5S.** Characterizing the expression patterns of ACE2 and TMPRSS2 in LG5 and LD6 cells. Total RNAs and proteins were extracted and analyzed by qRT-PCR and Western blot. (A) Bars indicate relative abundance of ACE2 and TMPRSS2 transcript levels normalized to RPL34 ribosomal protein RNA and expressed as fold-increase over control. (B) Representative immunoblots and quantitative analysis of ACE2 and TMPRSS2 protein levels in LD6 and LG5 cells. Data are presented as mean ± SD of three independent experiments. *$p < 0.05$, **$p < 0.005$. (C-F). NPC1 and ACE2 intracellular localization in LD6 and LG5 cells. Representative images of double immunofluorescence analyses performed by reacting cells with the following pairs of antibodies: α-ACE2 (red), α-LC3B (green) (C), α-ACE2 (green), α-LC3B (red) (D). Nuclei were stained with Hoechst 33342. Images are representative of at least three independent experiments. Scale bar: 50 μM.

**Supplementary Table 1.** List of oligonucleotides used as PCR primers.

| OLIGONUCLEOTIDE NAME | SEQUENCE 5'→ 3' |
|---|---|
| *H. sapiens* RPL34 fw | CCAGCGTTTGACATACCGAC |
| *H. sapiens* RPL34 rev | TGCTTTCCCAACCTTCTTGGT |
| *H. sapiens* NPC1 2$^{nd}$-3$^{rd}$ exons fw | TTCTGGCCCACCAAAACCATT |
| *H. sapiens* NPC1 2$^{nd}$-3$^{rd}$ exons rev | CTGCCGAACATCACAACAGAG |
| *C. sabeus* NPC1 5$^{th}$-6$^{th}$ exons fw | ACCATCACTCCCGTGTTTCA |
| *C. sabeus* NPC1 5$^{th}$-6$^{th}$ exons rev | GGGCCACAGACAATAGAGCA |
| *H. sapiens* ACE2 fw | TCACGATTGTTGGGACTCTGC |
| *H. sapiens* ACE2 rev | CCACCACCCCAACTATCTCTC |
| *C. sabeus* ACE2 fw | TCACGATTGTTGGGACTCTGC |
| *C. sabeus* ACE2 rev | CCACCACCCCAACTATCTCTC |
| *H. sapiens* TMPRSS2 fw | GGAGGACGAGAATCGGTGTG |
| *H. sapiens* TMPRSS2 rev | TCGTTCCAGTCGTCTTGGC |
| *C. sabeus* TMPRSS2 fw | ACTGCTGGATTTCTGGGTGG |
| *C. sabeus* TMPRSS2 rev | AGATCATGGCTGGTGTGACC |
| NPC1 4$^{th}$ exon fw | TGACATGTAGCCCTCGACAG |
| NPC1 4$^{th}$ exon rev | CTGTCCGACGTAGTATTGTAACTC |
| NPC1 4-5$^{th}$ exons fw | TGACATGTAGCCCTCGACAG |
| NPC1 4-5$^{th}$ exons rev | CATCCCGGCAGGCATTGTA |

### Fig1s

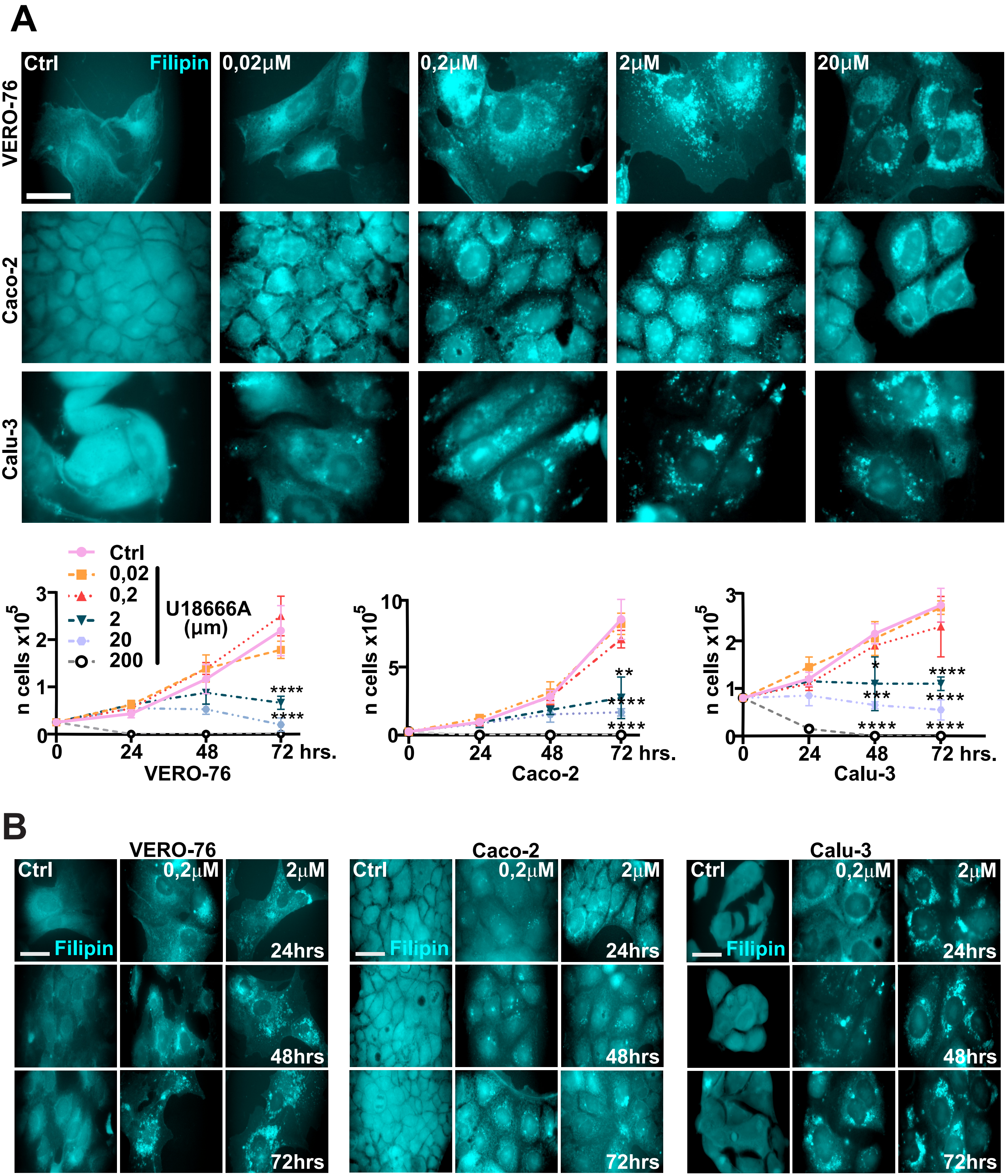

A
Ctrl
Filipin
0,02μM
0,2μM
2μM
20μM
VERO-76
Caco-2
Calu-3
Ctrl
0,02
0,2
2
20
200
U18666A
(μm)
n cells x10^5
0
1
2
3
0
24
48
72 hrs.
VERO-76
****
****
n cells x10^5
0
5
10
0
24
48
72 hrs.
Caco-2
**
****
****
n cells x10^5
0
1
2
3
0
24
48
72 hrs.
Calu-3
*
***
****
****
****
****
B
VERO-76
Caco-2
Calu-3
Ctrl
0,2μM
2μM
Filipin
24hrs
48hrs
72hrs

### Fig2s

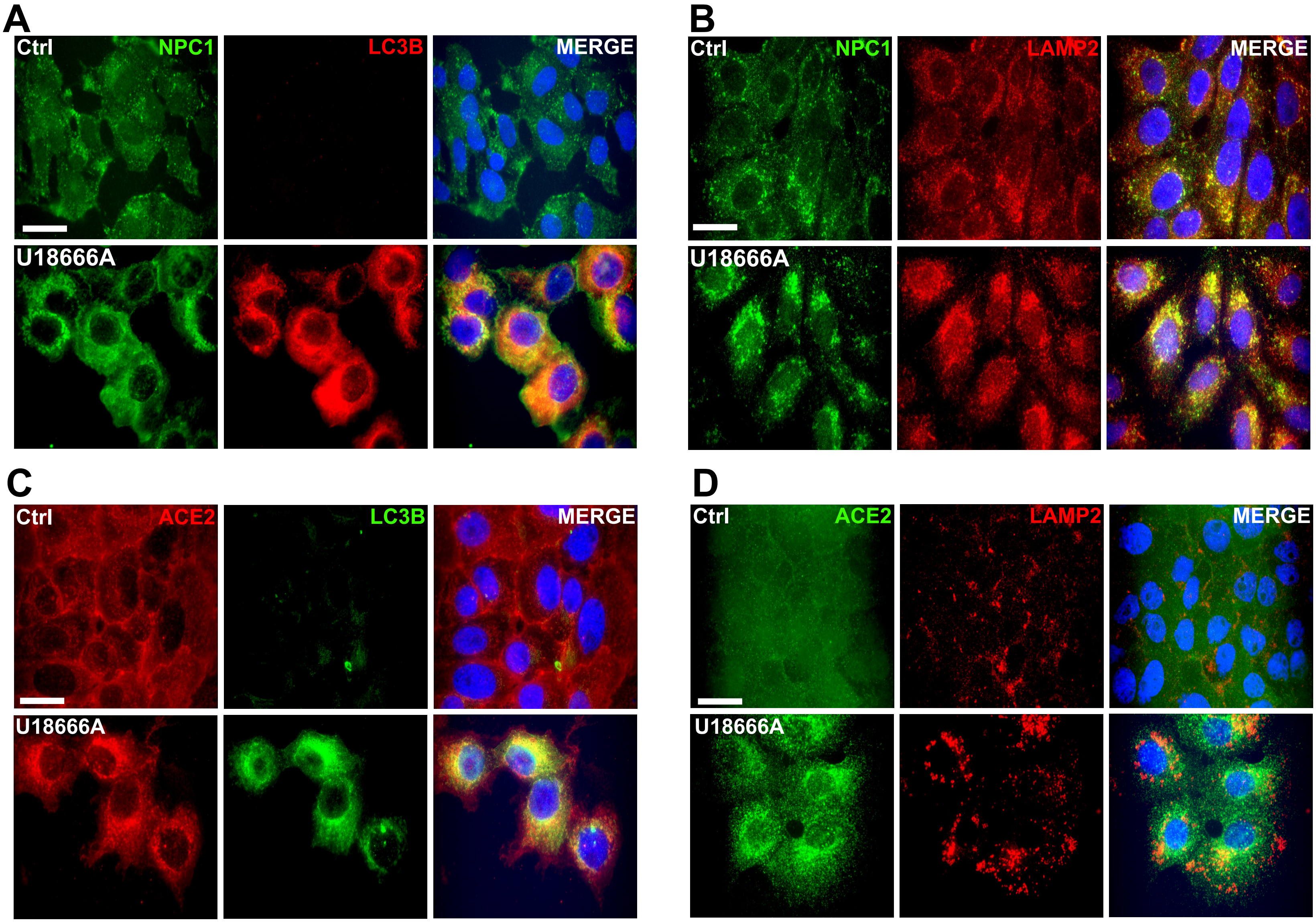

A
Ctrl
NPC1
LC3B
MERGE
U18666A
B
Ctrl
NPC1
LAMP2
MERGE
U18666A
C
Ctrl
ACE2
LC3B
MERGE
U18666A
D
Ctrl
ACE2
LAMP2
MERGE
U18666A

Fig3s

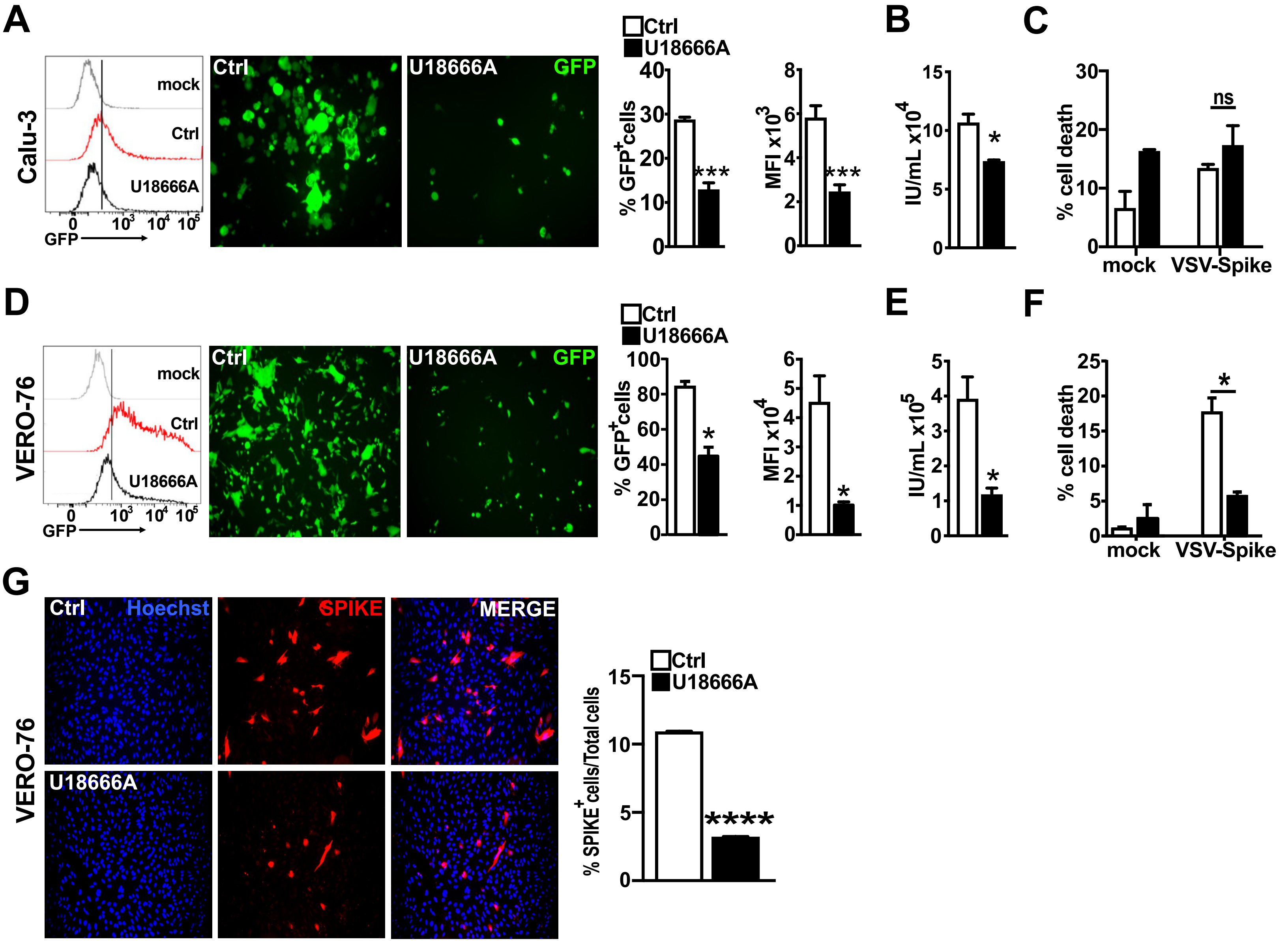

A
B
C
D
E
F
G
Calu-3
mock
Ctrl
U18666A
GFP
Ctrl
U18666A
GFP
% GFP+cells
MFI x10^3
IU/mL x10^4
% cell death
mock
VSV-Spike
ns
VERO-76
MFI x10^4
IU/mL x10^5
Hoechst
SPIKE
MERGE
% SPIKE+ cells/Total cells
***
****

### Fig4s

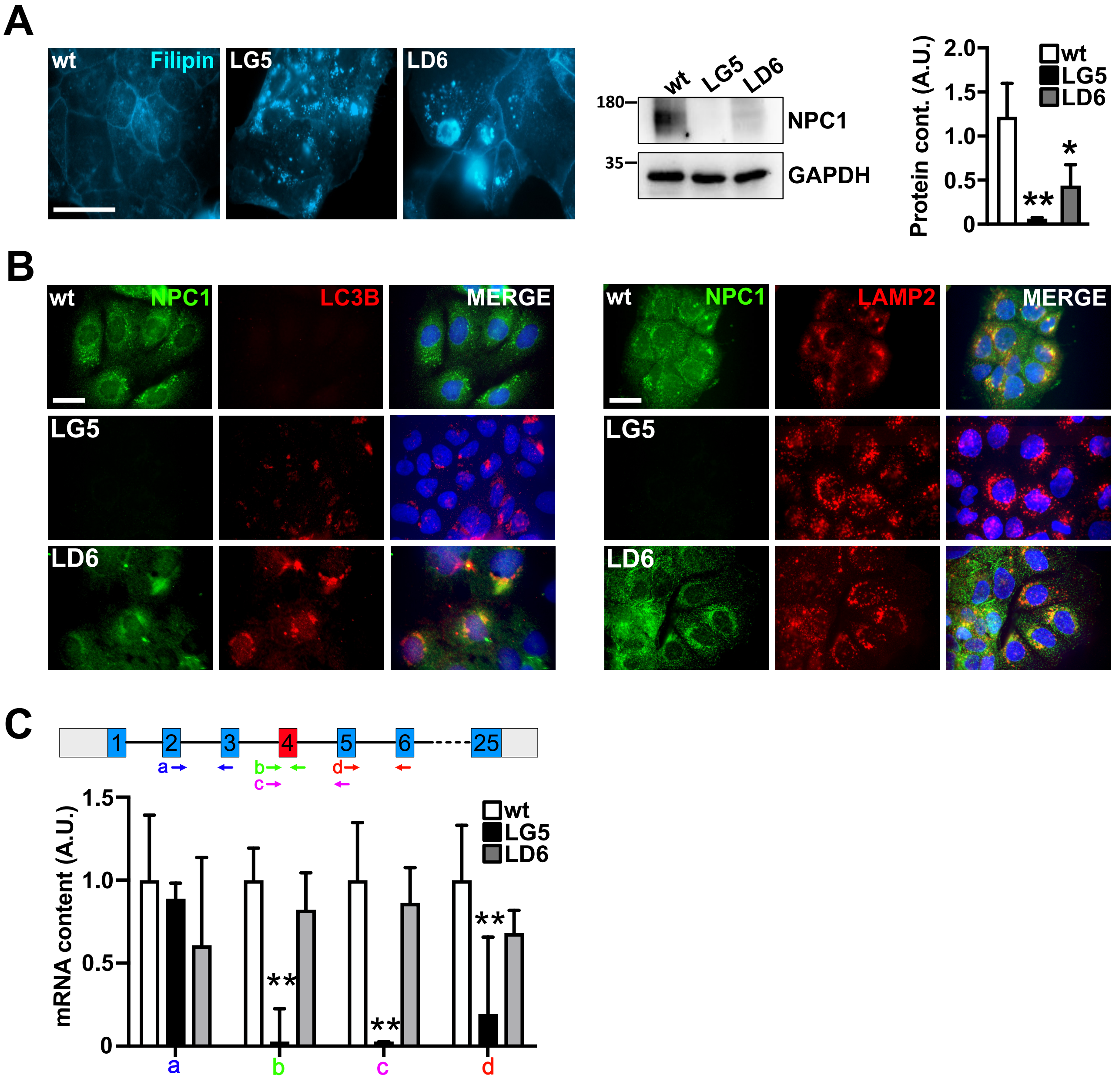

A
wt
Filipin
LG5
LD6
wt
LG5
LD6
180
35
NPC1
GAPDH
Protein cont. (A.U.)
2.0
1.5
1.0
0.5
0
wt
LG5
LD6
**
*
B
wt
NPC1
LC3B
MERGE
LG5
LD6
wt
NPC1
LAMP2
MERGE
LG5
LD6
C
1
2
3
4
5
6
25
a
b
c
d
mRNA content (A.U.)
1.5
1.0
0.5
0
wt
LG5
LD6
**
**
**
a
b
c
d

### Fig 5s

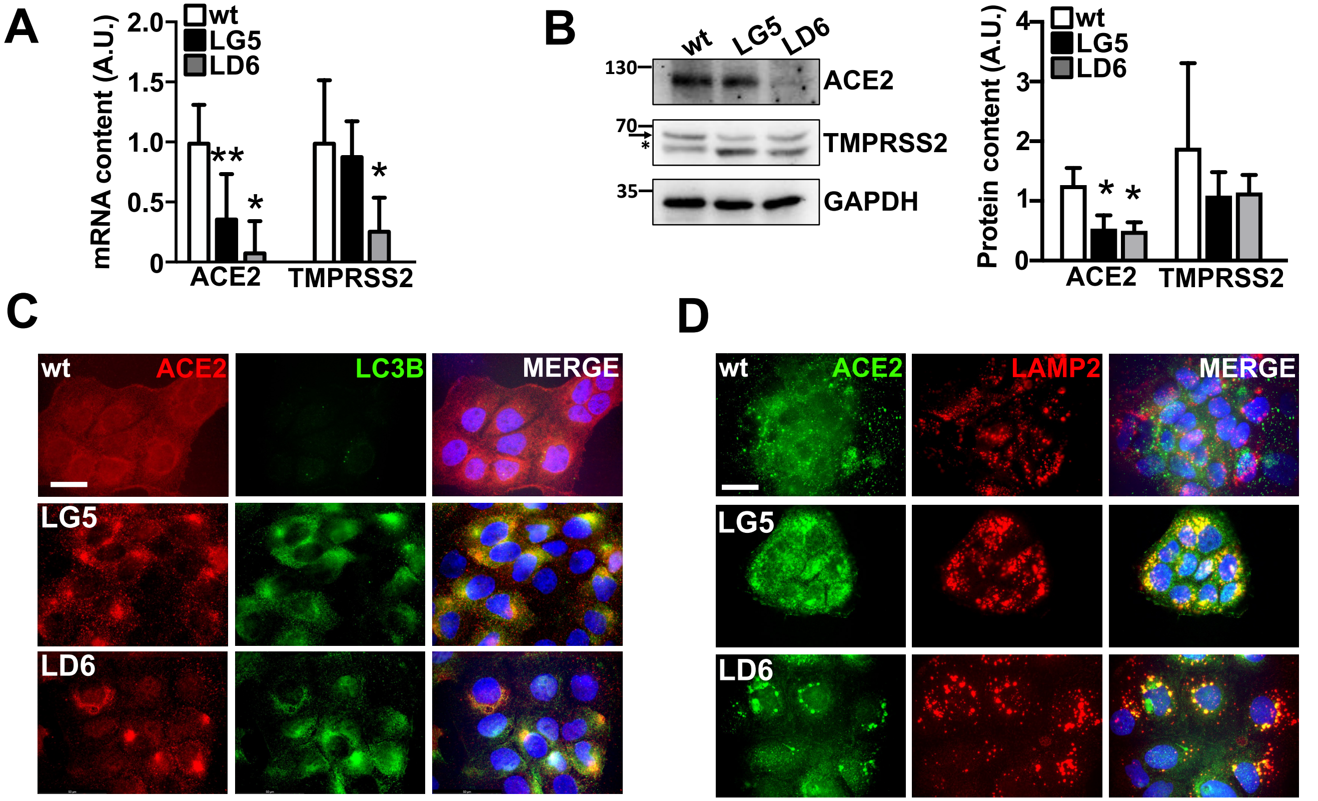

A
mRNA content (A.U.)
2.0
1.5
1.0
0.5
0
wt
LG5
LD6
**
*
*
ACE2
TMPRSS2
B
wt
LG5
LD6
130
70
35
ACE2
TMPRSS2
GAPDH
Protein content (A.U.)
4
3
2
1
0
ACE2
TMPRSS2
C
wt
ACE2
LC3B
MERGE
LG5
LD6
D
wt
ACE2
LAMP2
MERGE
LG5
LD6
